# Finding Expression of MUC1 and MUC4 in the Respiratory System of the Iraqi Common Quail (*Coturnix coturnix*)

**DOI:** 10.1101/2023.09.01.555941

**Authors:** Nabeel Abd Murad AL-Mamoori, Hazem Almhanna, Abdulrazzaq B Kadhim, David Kilroy, Arun HS Kumar

**Affiliations:** Department of Anatomy and Histology, College of Veterinary Medicine, University of Al-Qadisiyah. Iraq; Stemcology, School of Veterinary Medicine, University College Dublin, Belfield, Dublin-04, Ireland

**Keywords:** Mucus, Neutral Mucins, Acidic Mucins, MUC1, MUC4

## Abstract

**Background:** This study focused on the major components of mucus, known as mucins, within the mucosal epithelium of the respiratory system in Iraqi Common Quail. Six quail were utilized in accordance with animal ethics guidelines from the College of Veterinary Medicine at the University of Al-Qadisiyah. Histological analysis, utilizing H&E staining, aimed to identify key respiratory system structures. PAS plus Alcian blue stains were employed to identify specific carbohydrates in the trachea, bronchi, and lungs. RT-qPCR was used to assess the gene expression levels of MUC1 and MUC4.

**Results:** The trachea and bronchi encompassed four distinct layers: tunica mucosa, tunica submucosa, hyaline cartilage, and tunica adventitia. The mucosa consisted of pseudostratified epithelium that transitioned into simple columnar cells toward the primary and secondary bronchioles. This transition further progressed into simple cuboidal and squamous epithelium at smaller tertiary branches of the secondary bronchioles. Notably, the bronchial tunica submucosa was thinner than the trachea. While hyaline cartilage was prominently present in the trachea, it became fragmented in the bronchi and diminished towards the lungs and secondary bronchioles. Lung tissue was characterized by numerous lobules housing alveoli connected to alveolar ducts and sacs, alongside an intricate network of blood vessels. The respiratory tissues, including the trachea, bronchi, and lungs, exhibited a strong affinity for PAS-combined Alcian blue stains. This confirmed the substantial presence of both acidic and neutral mucins within the epithelial cells and glands. The trachea demonstrated significantly elevated levels of acidic mucins and a concentrated amount? of neutral mucins. Transcriptome analysis indicated the expression of both MUC1 and MUC4 genes. Importantly, MUC4 expression surpassed that of MUC1 in the trachea, bronchi, and lungs.

**Conclusion:** This study highlights the similarity of histological structures in the trachea, bronchi, and lungs of quail to typical avian species. Moreover, it underscored the substantial presence of both acidic and neutral mucins, with MUC4 being the predominant mucin, potentially playing a pivotal role in regulating mucosal barrier functions and interacting with pathogens. Nonetheless, further investigation is warranted to elucidate MUC4’s role in respiratory epithelial cells.

## Introduction

Mucins are glycoproteins composed of complex and intricate polymer structures which are notable for their substantial molecular size [1, 2]. These polymers are formed through the linkage of various amino acids, contributing to the creation of hydrogen bonds and distinct domains. Such molecular characteristics afford mucins a diverse range of interactions with both external and internal cellular elements in the body. These interactions are facilitated by a range of properties, including hydrophilic and hydrophobic traits, an array of sugar molecules, hydrogen bonding, and electrostatic interactions[3, 4]. Within the mucin family, numerous members are classified based on their specific expression locations in the body. This classification includes secreted mucins, which can be further categorized into gel-forming mucins and non-gel-forming (soluble) mucins [5, 6]. Additionally, there are transmembrane mucins (membrane-bound), which form part of the epithelial surface of cells [7, 8].

Mucins are expressed in various tissues, encompassing the digestive, respiratory and nervous systems, as well as the male and female reproductive tracts [9-13]. Mucin, as a pivotal constituent of mucus, assumes a crucial role in creating a physical barrier among epithelial cells. This barrier acts as a defense, impeding the adherence and localization of various infectious agents like viruses, bacteria, and parasites [14, 15]. This is particularly noteworthy in the avian respiratory system, where mucin expression helps birds guard against diverse infectious diseases [16, 17], offering a degree of protection [18]. Numerous pathogens invade the avian respiratory system, often leading to the increased secretion of mucins within mucus [19]. Unfortunately, this augmented mucus secretion can obstruct the airways, resulting in symptoms such as sneezing and coughing throughout the course of the disease [20-23]. Mucin 1 (MUC1), characterized by its high degree of glycosylation and substantial molecular weight [24, 25], finds its expression in the basal cell epithelium of the trachea and as a soluble form within the airways of the respiratory system [26]. Consequently, the levels of MUC1 could serve as an indicator for various diseases, offering insights into disease severity within the respiratory system [27, 28]. Furthermore, Mucin 4 (MUC4) which is a highly glycosylated glycoprotein, is characterized by its substantial molecular weight [29, 30]. This protein finds its expression on the apical surface of epithelial cells, playing a pivotal role in maintaining the integrity of the external environment within the respiratory system. It contributes significantly to the constitution of the mucosal barrier on the epithelium [31, 32]. Consequently, any aberrant expression of MUC4 could potentially serve as a valuable biomarker for a range of diseases [33, 34]. Aligned with this prior research, this study aimed to quantify the mucin types and assess the expression levels of both MUC1 and MUC4 in distinct respiratory system regions of Iraqi Common Quail. This comprehensive investigation aims to yield more expansive outcomes, contributing to a deeper understanding of mucin expression for various applications in research and serving as a foundation for future studies.

## Materials and Methods

### Samples and Design of Study

Six Iraqi common quail birds were selected as subjects for this study. These birds were in a healthy condition and were handled following the ethical guidelines outlined by the College of Veterinary Medicine at the University of Al-Qadisiyah, Iraq (Approval P.G. No. 1890). The Quail were euthanized by cervical dislocation. Subsequently, their internal organs were dissected, and samples of the trachea and lungs were collected for further analysis. Some of these samples were preserved in Trizole for subsequent Quantitative Real-Time PCR (qPCR) studies, while others were fixed in 10% formalin for histological examination.

### Histological study

The specimens of the trachea, bronchi, and lungs were directly collected immediately after the birds were euthanized. These collected samples were then fixed in a 10% formalin solution (at a ratio of 1:10) for a period of 48 hours [35-37]. Following the fixation process, the tissue underwent preparation for histological analysis. This involved the application of staining techniques, including Hematoxylin and Eosin (H&E) staining, as well as a combination of Periodic Acid-Schiff reagent (PAS) and Alcian blue, following the established protocol [38]. Briefly, the samples were initially immersed in tap water for a duration of two hours. Subsequently, the samples underwent a series of alcohol-based immersions for dehydration, with each immersions lasting for two hours. Following this, the samples were subjected to two cycles of xylene immersion, with each cycle lasting for two hours. The next step involved soaking the samples in paraffin wax (at 56°C) for two hours, after which the tissue blocks were prepared. Consequently, tissue sections were sliced onto slides using a microtome. These slides were subjected to staining processes, utilizing H&E as well as a combination of Alcian blue and PAS. A light microscope (Olympus, Japan) was employed to examine the slides, capturing images of the tissue at varying magnifications (4x, 10x, 20x, and 40x).

### RNA Extraction

The RNA expression profiles of the trachea, bronchi, and lung tissues of Iraqi common Quail were extracted following the instructions of the Accuzol® reagent kit (Bioneer, Korea). A total of 150 mg was precisely weighed and collected from each section of the trachea, bronchi, and lung tissues, with each segment being carefully placed into a 1.5ml Eppendorf tube. The specimens were then treated with 200μl of chloroform (Sigma-Aldrich, USA), mixed thoroughly, and subsequently incubated on ice for a duration of 5 minutes. Following the incubation, the specimens underwent centrifugation at 10,000 rpm and 4°C for a period of 15 minutes, with only the supernatant being retained. To this supernatant, an equal volume (1:1) of isopropanol (USA) was added, ensuring thorough mixing, and the mixture was incubated once again at 4°C for 10 minutes. Subsequently, the specimens were subjected to centrifugation at 10,000 rpm and 4°C for 10 minutes, and the supernatant was discarded, leaving the pellets intact. Next, the pellets were exposed to 1ml of 80% ethanol (Sigma-Aldrich, USA) and mixed once more using a vortex. The specimens were then subjected to centrifugation at 10,000 rpm and 4°C for 10 minutes, followed by the removal of the supernatant and allowing the pellets to air-dry. Finally, the RNA pellets were reconstituted in 50μl of DEPC water (Sigma-Aldrich, USA), and the resulting solution was stored at a temperature of - 20°C. Additionally, the concentration of RNA was assessed for each sample using a Nanodrop spectrophotometer (THERMO, USA).

### cDNA synthesis

cDNA of extraction RNA was made using a commercial kit (Promega Company, USA), and RNA of samples was incubated with DNase I enzyme for two hours to remove any remains of DNA contamination. After that, purified RNA was translated into cDNA using DiaStar™ OneStep RT-PCR Kit (China) according to the instructions of the kit and thermocycler conditions. Finally, normalization of cDNA concentrations was applied for samples and kept at -20°C for the next steps.

### Quantitative reverse transcription polymerase chain reaction (RT-qPCR)

RT-qPCR was employed to find the levels of quantification of mucin 1 and 4 mRNA transcripts according to protocol of the Real-Time PCR system (BioRad./ USA)[39]. Primers for this study were made by Scientific Researcher Co. Ltd in Iraq and included Coturnix japonica glyceraldehyde-3-phosphate dehydrogenase (GAPDH housekeeping gene), and Coturnix japonica MUC1-like (LOC107317569) (MUC1), and Coturnix japonica mucin 4, cell surface associated (MUC4), listed in table (1):

**Table 1:**
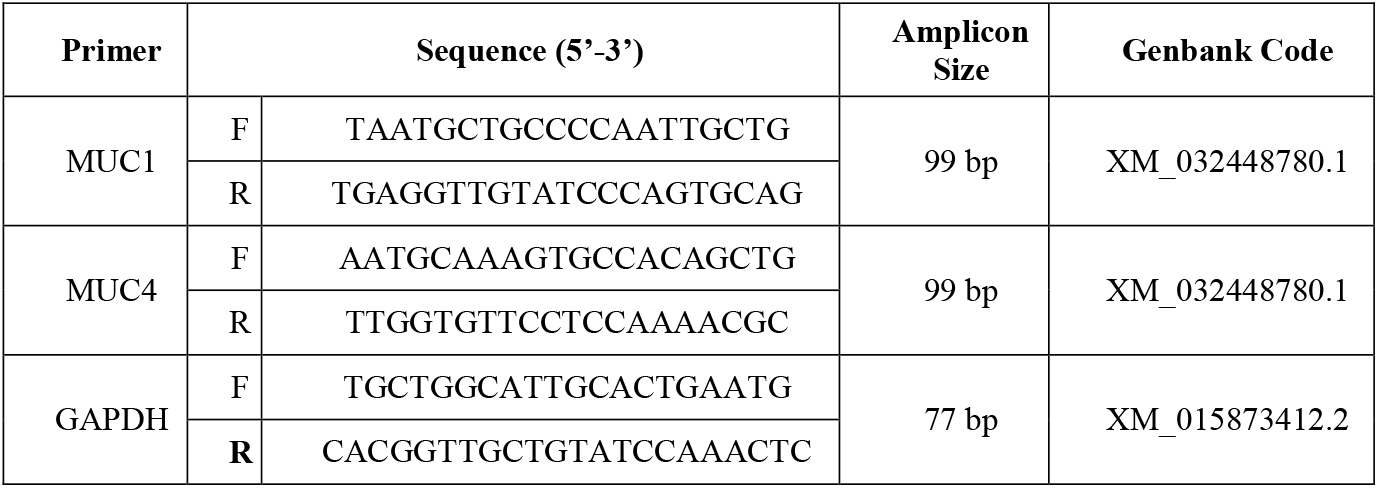
The Primer sequence used for the detection of MUC1, Muc4 and GAPDH.

The SYBR Green dye qPCR master mix and kit (AccuPowerTM 2X Green Star qPCR master mix kit, Bioneer, Korea) were employed to facilitate the detection, amplification, and normalization of gene expression levels. This encompassed the analysis of the GAPDH housekeeping gene, as well as the MUC1 and MUC4 genes. The qPCR thermocycler conditions were established as follows: The initial denaturation phase was set at 50°C for an hour, involving a single cycle. The subsequent cycle commenced with denaturation at 95°C for 20 seconds, followed by annealing and extension detection (scanning) at 60°C for 30 seconds. This cycle was repeated with a duration of 45 seconds. Finally, the melting temperature was maintained within the range of 60°C to 95°C, with a duration of 0.5 seconds. This cycle was executed only once.

### Statistical Analysis

The quantities of neutral and acidic mucins in the trachea, bronchi, and lungs were measured using Image J. The means of mucin quantities were then analyzed. Additionally, the raw data of gene expression for the MUC1 and MUC4 genes in the same tissues were detected through RT-qPCR and analysed using the 2^ΔCT method [40, 41]. For statistical analysis, IBM SPSS Statistics 23.0 (SPSS) was utilized. A two-way ANOVA was applied to analyse the mucin quantities, while a one-way ANOVA was employed to analyse the RT-qPCR data. Significance was considered when the p-value≤0.05.

### Results

A histological analysis of the trachea, bronchi, and lungs of Iraqi common Quail was performed using H&E staining as well as a combination of PAS and Alcian blue stains. The trachea was composed of three layers, including the tunica mucosa, tunica submucosa reinforced by cartilage, and the outermost tunica adventitia. The tunica mucosa was lined by ciliated pseudostratified columnar epithelium with goblet cells interspersed between the epithelial cells. These goblet cells were more prominent in the distal portions of the trachea. This layer was supported by a thin lamina propria that interacted with the submucosa. The submucosa, the largest layer, was occupied by thick hyaline cartilage and surrounded by loose connective tissue containing collagen fibers, fibrocytes, lymph cells, adipose tissue, small blood vessels, and additional spaces. The hyaline cartilage formed complete rings and contained binucleated chondrocytes. The tunica adventitia consisted of connective tissue along with smooth muscle fibers oriented in different directions. Staining the trachea with the PAS/Alcian blue combination showed that the epithelial layer was stained more with PAS and less with Alcian blue. Goblet cells, tracheal glands, and hyaline cartilage stained strongly with Alcian blue (Figure 1). The analysis of mucin quantities in the trachea revealed that acidic and neutral mucins were both highly expressed in epithelial goblet cells and tracheal glands. Expression of acidic and neutral mucin was similar in the trachea and bronchi; however, in the lung tissue the expression of acidic mucin was significantly (p<0.01) higher than expression of neutral mucin (Figure 1). While the expression of neutral mucin was similar across trachea, bronchi and lungs, we did observe differences in the expression pattern of acidic mucins across these three tissues. The expression of acidic mucins was highest in the trachea and lowest in the lungs (Figure 1).

**Figure 1:**
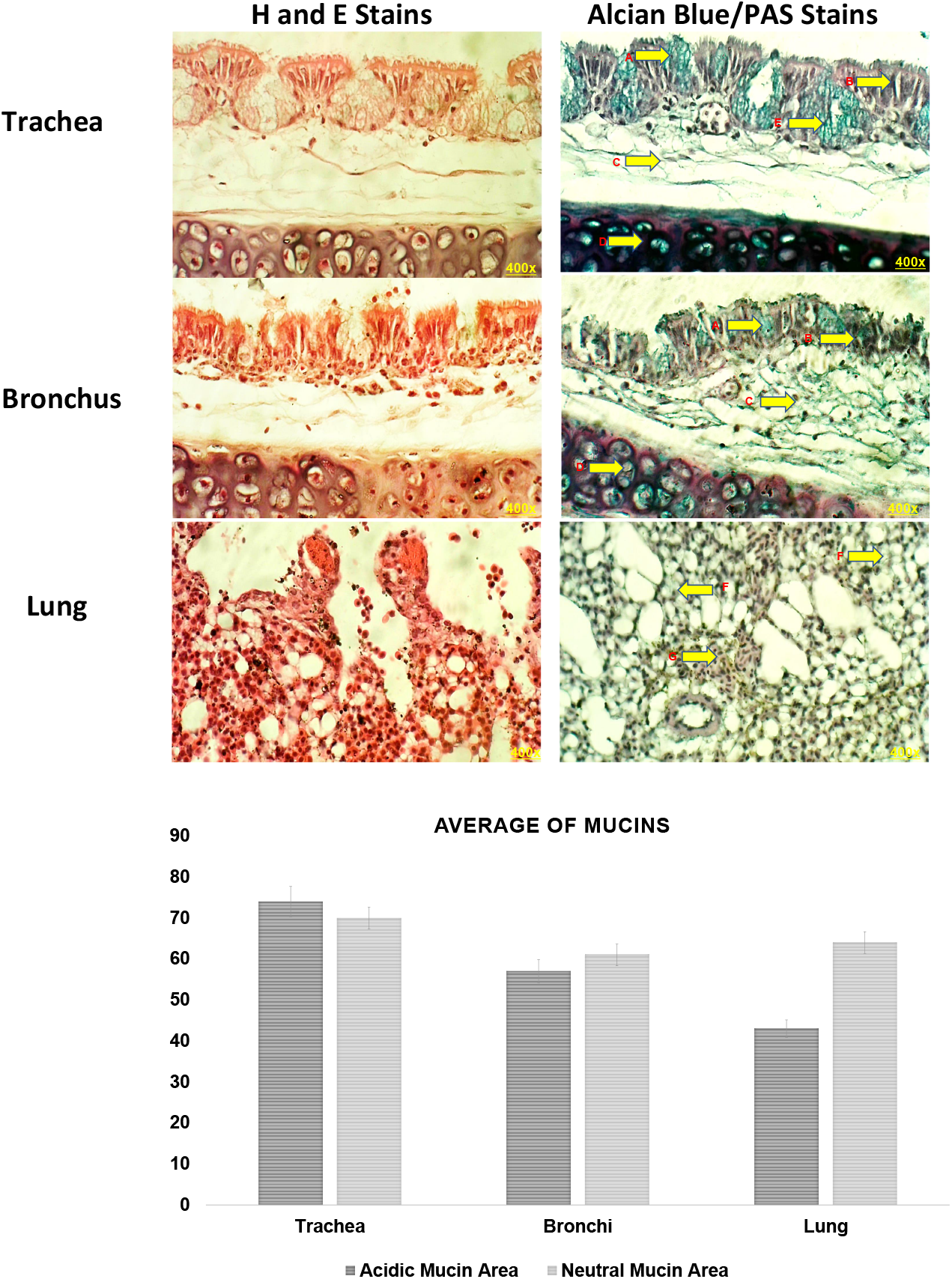
Representative section of quail trachea, bronchi and lungs stained with H&E stain or alcian blue/PAS stain. The bar graph shows quantification of the acidic and neutral mucin staining in quail trachea, bronchi and lungs. Data is represented as Mean +/-SEM (n=3). * p<0.05, ** p<0.01, *** p<0.001.

The bronchi exhibited similar structural patterns to the trachea. However, the cartilage was interrupted and gradually disappeared as they extended towards the lungs. The epithelium of the bronchi was lined by pseudostratified columnar ciliated epithelium. Variable numbers of mucous glands and lymphoid tissue were present in the lamina propria. The bronchi gave rise to numerous branches that extended into the lung lobules. These branches divided into primary, secondary, and tertiary bronchioles. The epithelium of these bronchioles was similar to that of the bronchi, but they had fewer or no cartilaginous rings (Figure 1). As these bronchioles extended into the lungs, they transformed into parabronchi and the epithelium changed from simple cuboidal to squamous epithelium, which continued into the atria and infundibula. The PAS stain highlighted the epithelium of the bronchi and their smaller branches, while Alcian blue stained the goblet cells, mucosal glands, and cartilage. The quantity of neutral mucins in the bronchi exhibited a relative increase compared to acidic mucins; however this was not statistically significant (p>0.05). Additionally, the lung revealed the presence of alveolar ducts, air sacs, spaces, and septa that clearly divided the lung into lobules. These structures were predominantly stained with PAS and to a lesser extent with Alcian blue (Figure 1). The PAS and Alcian blue stains also highlighted the epithelium of the small branches of the bronchioles within the lungs. The lung tissue exhibited a significantly higher proportion of neutral mucin in comparison to acidic mucin (p<0.01) and the lung showed least expression of acidic mucins compared to the bronchi and trachea (p<0.05 and 0.01 respectively).

The RT-PCR amplification plots of the MUC1 and MUC4 genes in the tissues of Iraqi common Quail displayed distinctive Ct cycle numbers, ranging between 27 and 34 (Figure 2). The RT-qPCR analysis of housekeeping genes and MUC1 and MUC4 genes exhibited high specificity and consistent curve amplifications with distinct melting peaks. These melting peaks for the housekeeping gene, MUC1, and MUC4 were in the ranges of 75-85°C, 84-89°C, and 82-89°C respectively (Figures 2). The RT-qPCR amplifications of mucin genes indicated that the mRNA of MUC1 and MUC4 were indeed expressed in the trachea, bronchi, and lungs (Figures 2), as presented in Tables 2 and 3. Although the concentration of MUC4 was notably higher in the trachea, bronchi, and lungs compared to MUC1, this difference was not statistically significant (Figure 2, Tables 2 and 3). Notably, MUC4 exhibited elevated expression levels in the bronchi, surpassing those in the trachea and lungs. Additionally, MUC1 expression levels were relatively similar in the trachea, bronchi and lungs.

**Table 2:** Values of gene expression and housekeeping gene of MUC1 which were analysed using 2^ΔCT method.

| No. | Tissue | CT (MUC1) | CT (GAPDH) | $\Delta CT$ | Fold change ($2^{\Delta\Delta CT}$) | Mean |
| --- | --- | --- | --- | --- | --- | --- |
| 1 | Trachea | 30.54 | 30.93 | 0.39 | 1.31 | 1.90 |
| 2 | Trachea | 33.02 | 31.35 | -1.67 | 0.31 |  |
| 3 | Trachea | 28.21 | 30.24 | 2.03 | 4.08 |  |
| 4 | Bronchi | 31.17 | 30.36 | -0.81 | 0.57 | 5.30 |
| 5 | Bronchi | 34.47 | 30.99 | -3.48 | 0.09 |  |
| 6 | Bronchi | 26.79 | 30.72 | 3.93 | 15.24 |  |
| 7 | Lung | 28.77 | 30.12 | 1.35 | 2.55 | 5.29 |
| 8 | Lung | 36.06 | 31.83 | -4.23 | 0.05 |  |
| 9 | Lung | 26.79 | 30.52 | 3.73 | 13.27 |  |

**Table (3):**
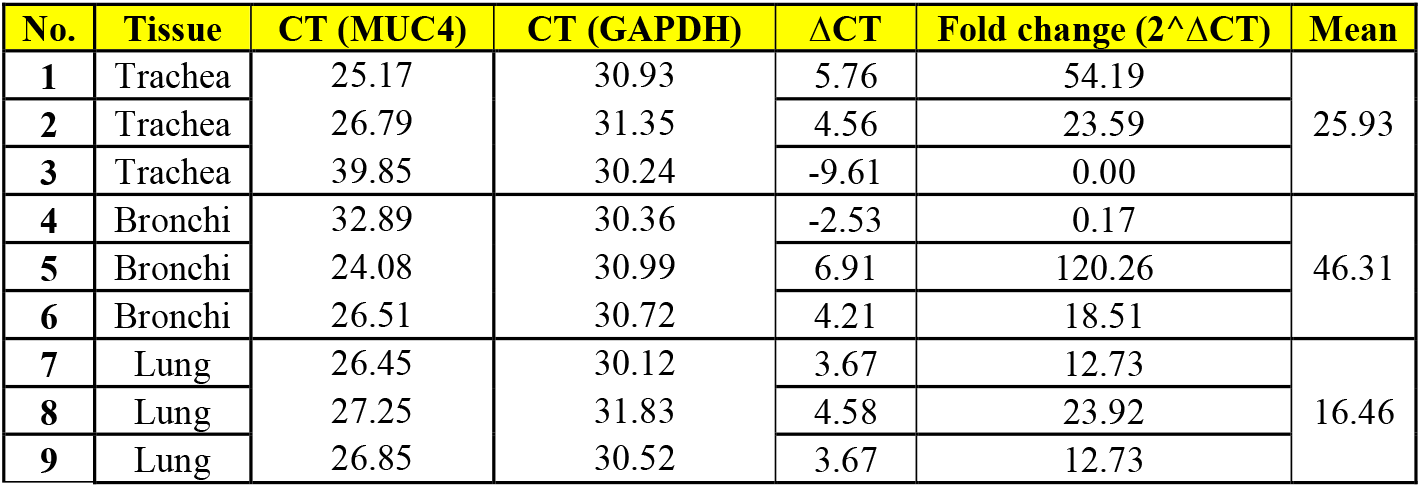
Values of gene expression and housekeeping gene of MUC4 which were analysed using 2^ΔCT method.

**Figure 2:**
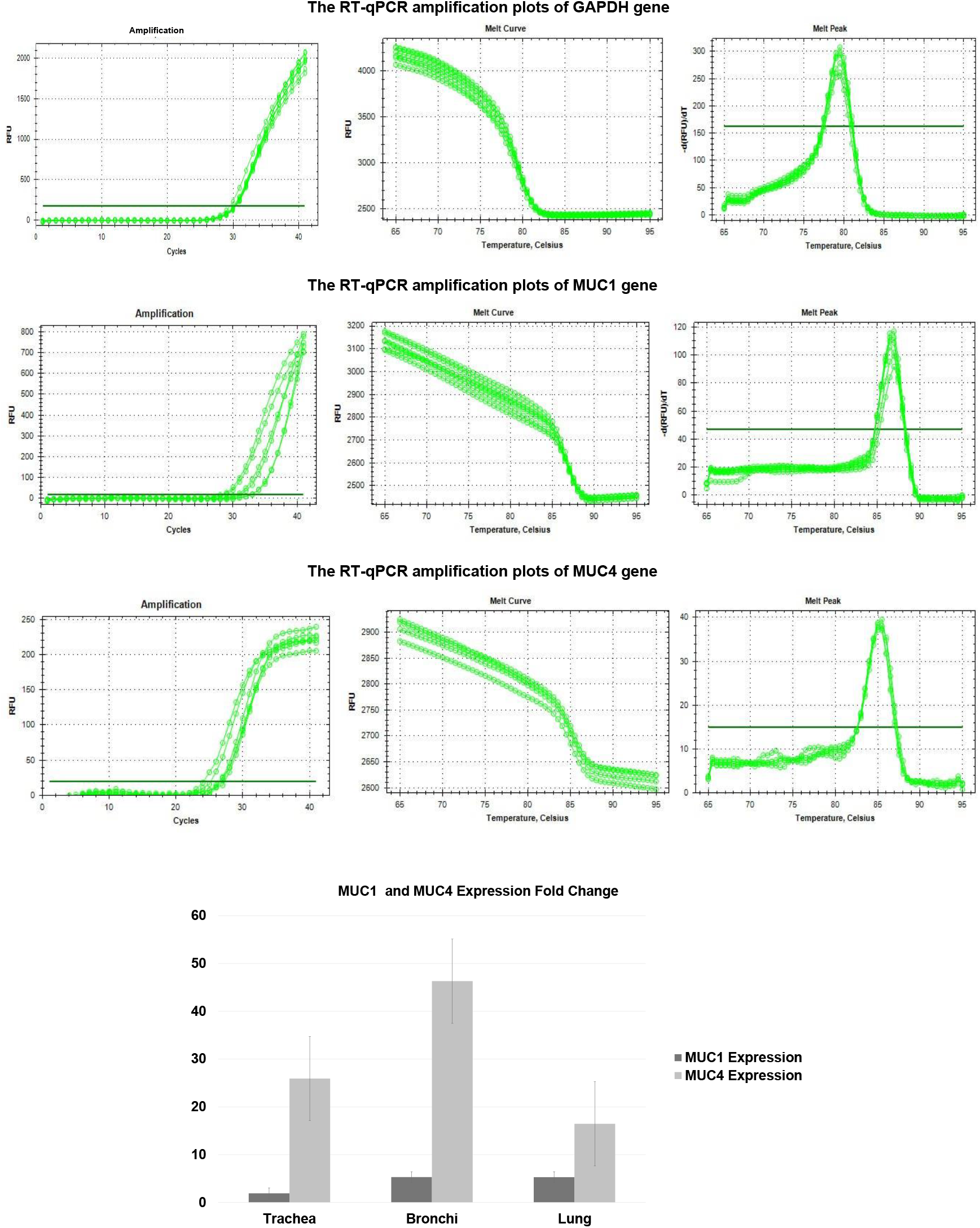
Quantification of the MUC 1 and 4 transcripts in quail trachea, bronchi and lung tissue using RT-qPCR. The expression of MUC 1 and 4 was normalised to expression of GAPDH.

## Discussion

Previous studies have reported that mucins (highly glycosylated glycoproteins consisting of different members) are expressed on epithelial cells throughout the body, including the respiratory, digestive, and urogenital tracts, as well as in accessory sex glands of the male reproductive system, mammary glands, and salivary glands [13, 42, 43]. Therefore, our study which focused on various types of mucins, particularly MUC1 and MUC4, in the respiratory system of the Iraqi common quail is consistent with these previous reports. Studies conducted on Japanese Quail indicated the presence of variable mucin types, including both neutral and acidic mucins, which exhibited changes corresponding to the post-hatching period of the Quail [44]. These findings are consistent with our results, which confirm the presence of a mixture of mucins, with acidic mucins being predominant over neutral mucins, and this predominance diminishes towards the lungs. This suggests that acidic mucins may play a role in maintaining moisture in the trachea, preventing blockages, and serving as a defence against bacterial infections. In the case of Iraqi common Quail, MUC1 and MUC4 have been found to be expressed in the digestive system [45]. The respiratory airway contains a variety of mucins, including gel-forming mucins (MUC5AC, MUC5B) and transmembrane mucins (MUC1, MUC4, MUC16, MUC20) [46]. Moreover, evidence suggests the expression of additional MUC genes such as MUC2, MUC7, MUC8, MUC11, MUC13, MUC15, and MUC19 [47]. Notably, three cell surface-associated proteins—MUC1, MUC4, and MUC16—have been identified in the respiratory tract. These proteins are integral to cellular transduction pathways and transcription, regulating the diverse functions of the airway epithelium [43, 48, 49]. These findings align with our results, which identify the presence of MUC1 and MUC4 in the trachea, bronchi, and lungs, although MUC6 was not evaluated in our study and it may merit assessing this mucin type in future studies.

MUC1 and MUC4 are large transmembrane glycoproteins found in respiratory epithelial cells. They play a crucial role in protecting against extracellular secretions and infestations from parasites, bacteria, and viruses [50, 51]. This aligns with our findings, which suggest that the expression of MUC1 and MUC4 in the trachea and lung tissues of Quail contributes to regulating immune responses and safeguarding epithelial cells from infectious lesions. These proteins contribute to forming the mucosal barrier that covers the respiratory epithelium and lung tissue, in addition to aiding in gas exchange between alveolar sacs and the capillary bed. In our study, we found that MUC4 was highly expressed in the trachea, bronchi, and lungs, in contrast to MUC1 expression. This aligns with previous research [45] that has indicated that MUC4 has larger ligand binding domains compared to MUC1. This observation supports the idea that MUC4 could play a central role in constituting the primary mucosal barrier of the airway epithelium, defending against pathogenic infections, and regulating the glycosylation and sulfation of other mucins by controlling the secretion of sulphide from epithelial cells. The mucosal barrier of the respiratory system is also rich in mucins with diverse sialic acids and other oligo- and polysaccharides [52]. Sialic acids and these other molecules have a high affinity for various viruses, contributing to the infectivity of pathogens such as coronaviruses, influenza, parainfluenza, and rotaviruses [53, 54]. In Quail, sialic acids carry acid receptors that facilitate strong binding of avian and human influenza viruses [55]. These findings, combined with our own study, suggest that MUC1 and MUC4, as transmembrane proteins in epithelial cells, likely play a key role in defending the respiratory system and detecting viral infections. Furthermore, in cystic fibrosis (CF) which is a hereditary chronic disease affecting the respiratory and digestive systems consequent to disruptions in ion and water movement across epithelial cells, leading to alterations in mucus components, the functional role of MUC1 and MUC4 may be vital [56, 57]. This is a crucial insight into the roles of MUC1 and MUC4 in the respiratory system influencing protein formation and cellular processes in CF warrants further research. The extensive binding sites of MUC4 suggest that it may bind to numerous soluble proteins, soluble mucins, and sugars, potentially explaining its higher concentration compared to MUC1. MUC4’s functions likely regulate cellular signal reactions in respiratory epithelial cells, maintaining typical mucus patterns in the airways according to extracellular conditions and diseases.

In conclusion, our study has identified the presence of both acidic and neutral mucins throughout the trachea, bronchi, and lungs of Quail. Additionally, we have confirmed the expression pattern of MUC1 and MUC4 in these tissues. MUC4 emerges as a major member of the mucin family, potentially regulating immune responses and serving as a crucial mucosal barrier in the airways. Any disruptions in MUC4 expression might be linked to the onset of various diseases, warranting further investigation.

## Notes

### Competing Interest Statement

The authors have declared no competing interest.

